# *NONEXPRESSOR OF PATHOGENESIS-RELATED GENES* control Huanglongbing tolerance by regulating immune balance in citrus plants

**DOI:** 10.1101/2024.03.18.585579

**Authors:** Poulami Sarkar, Choaa El-Mohtar, Donielle Turner, Stacy Welker, Cecile J. Robertson, Vladimir Orbovic, Zhonglin Mou, Amit Levy

**Affiliations:** Citrus Research and Education Center, Lake Alfred, University of Florida, FL 33850; Department of Microbiology and Cell Sciences, University of Florida, Gainesville, FL 32603; Department of Plant Pathology, University of Florida, Gainesville, FL 32602

**Keywords:** citrus, Huanglongbing, *Candidatus* liberibacter asiaticus, systemic acquired resistance, *NONEXPRESSOR OF PATHOGENESIS-RELATED GENES*, callose, reactive oxygen species, phloem plugging, tolerance

## Abstract

Huanglongbing (HLB) is a devastating citrus disease caused by the phloem-resident bacterial pathogen *Candidatus* liberibacter asiaticus (*C*Las). *C*Las infection of susceptible varieties triggers unbalanced immune responses, leading to overaccumulation of callose and reactive oxygen species (ROS), which in turn causes phloem plugging and HLB symptom development. Interestingly, some citrus relatives exhibit little or no symptoms in the presence of *C*Las, a phenomenon termed HLB tolerance. Moreover, overexpression of the *Arabidopsis thaliana NPR1* (*AtNPR1*) gene in susceptible varieties has been shown to confer robust HLB tolerance. However, the mechanisms underlying HLB tolerance remain enigmatic. Here, we show that overexpression of *AtNPR1* suppresses *C*Las- and *Pseudomonas syringae* pv. *maculicola* ES4326 (*Psm*)-induced overaccumulation of callose and ROS in citrus and *Arabidopsis*, respectively. Importantly, we found that knocking out of the *Arabidopsis* negative immune regulators, *AtNPR3* and *AtNPR4*, and silencing of their *Citrus sinensis* ortholog *CsNPR3*, similarly suppress *Psm*- and *C*Las-induced callose and ROS overaccumulation, respectively, and that silencing of *CsNPR3* also enhances HLB tolerance. These results reveal a conserved role of the *NPR1*/*NPR3*/*NPR4*-mediated signaling pathway in regulating plant immune balances and provide mechanistic support for overexpression of *AtNPR1* or silencing of *AtNPR3/AtNPR4* orthologs in citrus as a long-term solution to the HLB disease.

## Introduction

Huanglongbing (HLB) or citrus greening, caused by the bacterial pathogen *Candidatus* liberibacter asiaticus (*C*Las), is a devastating disease that afflicts citrus trees globally, leading to substantial production costs and declines in fruit quality. *C*Las infection triggers robust phloem immune responses characterized by excessive accumulation of reactive oxygen species (ROS) and increased deposition of callose. While a basal level of callose is transiently present in healthy sieve plates, callose can rapidly accumulate and block the sieve pores upon pathogen attack ^1^. ROS are also naturally produced during normal plant growth; however, their accumulation becomes excessive following *C*Las infection, leading to oxidative cell death ^2,3^. These responses trigger phloem collapse which obstructs the movement of photoassimilates source leaves to sink tissues ^4–8^. This obstruction is believed to cause the development of HLB symptoms including stunted growth, yellow and blotchy leaves, and lopsided fruits with aborted seeds ^9,10^. Current strategies to manage HLB symptom development and its spread mostly rely on foliar application and trunk injection of antimicrobials, with limited effectiveness and unknown physiological effects on the trees ^11–13^.

Comparative studies of HLB susceptible and tolerant citrus varieties have uncovered a potential role of immune responses, particularly systemic acquired resistance (SAR), in mediating HLB tolerance ^14,15^. SAR is a long-lasting broad-spectrum immune response induced by mobile signals produced at the primary infection site and requires the phytohormone salicylic acid (SA) ^16,17^. A key player in SA-mediated signaling is NONEXPRESSOR OF PATHOGENESIS-RELATED GENES1 (NPR1), which acts as an SA receptor ^18,19^. Upon SA-induced redox changes, NPR1 is converted from an oligomeric complex to monomers, which move into the nucleus where monomeric NPR1 interacts with TGA transcription factors to activate the transcription of defense genes including *PR* genes ^20–27^. The NPR1 paralogs, NPR3 and NPR4, are also SA receptors ^28^. NPR1 and NPR3/NPR4 play opposite roles in regulating SA-induced transcriptional changes ^29^. Interestingly, upon *C*Las infection, four *NPR1*-like genes were significantly upregulated in HLB-tolerant ‘Jackson’ grapefruit-like-hybrid trees and only one was upregulated in the closely related susceptible ‘Marsh’ grapefruit trees ^14^. Moreover, *NPR1*-like genes were not induced by *C*Las infection in susceptible ‘Valencia’ sweet orange ^30^. Additionally, overexpression of the *Arabidopsis thaliana NPR1* (*AtNPR1*) gene or knocking out the potato *NPR3* ortholog in microbial hairy roots significantly decreased the titers of *Candidatus* liberibacter solanacearum, which causes potato zebra chip, a disease similar to HLB ^31^. These results suggested a potential correlation between NPR1/NPR3-mediated signaling and HLB tolerance in citrus.

The importance of NPR1-mediated signaling in citrus disease resistance or tolerance has been underscored by multiple transgenic studies. Overexpression of *AtNPR1* or its citrus ortholog *CtNH1* (*Citrus NPR1 homolog 1*) in ‘Hamlin’ sweet orange and/or ‘Duncan’ grapefruit was initially found to enhance resistance to citrus canker ^32,33^. Subsequent intensive greenhouse screening identified independent transgenic lines that accumulate high levels of the AtNPR1 protein and only occasionally displayed mild HLB symptoms, and the progeny plants of these transgenic lines retained the same levels of HLB tolerance ^34^. Independently produced transgenic ‘Hamlin’ and ‘Valencia’ sweet orange expressing *AtNPR1* also exhibited reduced HLB disease severity under high disease pressure in the field ^35^. Although overexpression of *AtNPR1* in citrus has been shown to elevate the expression of a group of defense-related genes including *PR1* ^35,36^, the mechanisms preventing HLB disease symptom development remains unclear.

In this study, we characterized *C*Las-induced callose deposition and ROS accumulation in transgenic ‘Hamlin’ sweet orange and ‘Duncan’ grapefruit trees which overexpress the *AtNPR1* gene (hereafter *AtNPR1-OE*). We also examined the role of *Citrus sinensis NPR3* (*CsNPR3*), the closest citrus ortholog of *AtNPR3* and *AtNPR4* that encode two negative immune regulators ^37^ in HLB via *Citrus tristeza virus*-based RNA interference (CTV-RNAi) ^38^. Our results show that *AtNPR1* and *CsNPR3* oppositely regulate basal callose levels, *C*Las-induced callose and ROS accumulation, as well as HLB symptom development. Furthermore, we found that *AtNPR1* and *AtNPR3*/*AtNPR4* also oppositely regulate the bacterial pathogen *Pseudomonas syringae* pv. *maculicola* ES4326 (*Psm*)-induced callose and ROS accumulation in *Arabidopsis*. These results suggest a conserved function of *NPR1*/*NPR3*/*NPR4* in regulating plant immune balances and demonstrate that genetic manipulation of the signaling pathway mediated by *NPRs* in citrus is an effective approach to creating HLB tolerance.

## Results

### Callose deposition occurs early and precedes ROS accumulation in response to *C*Las infection

Overaccumulation of callose and ROS is believed to stimulate HLB symptom development ^4^. It has been shown that H_2_O_2_ and callose accumulate to significantly higher levels in young flushes produced by *C*Las-positive citrus plants than in those on healthy plants at 15- and 18-day post-bud initiation, respectively, indicating that *C*Las induces ROS production earlier than callose deposition in new flushes on *C*Las-positive plants ^4^. To determine the kinetics of *C*Las-induced callose and ROS accumulation in naïve healthy citrus plants, we inoculated ‘Hamlin’ (HAM) sweet orange and ‘Duncan’ (DUN) grapefruit with *C*Las-free (*C*Las-) and *C*Las-infected (*C*Las+) psyllids and analyzed callose deposition and ROS accumulation at 1-day and 14-day post-inoculation (dpi). Surprisingly, significant induction of callose deposition occurred within 1 dpi with *C*Las-infected psyllids and callose levels further increased at 14 dpi (Fig. 1a, b). In contrast, inoculation with *C*Las-free psyllids did not increase callose levels in the phloem of either the ‘Hamlin’ or ‘Duncan’ plants (Fig. 1a, b). Moreover, significant ROS accumulation, as measured by 3,3’-diaminobenzidine (DAB) staining, was detected at 14 dpi with *C*Las-infected psyllids (Supplementary Fig. S1; Fig. 1c, d). Inoculation with *C*Las-free psyllids did not cause significant ROS generation compared to the uninoculated controls. Taken together, these results indicate that *C*Las-triggered callose deposition happens considerably earlier than previously thought and is followed by ROS accumulation.

**Fig. 1.**
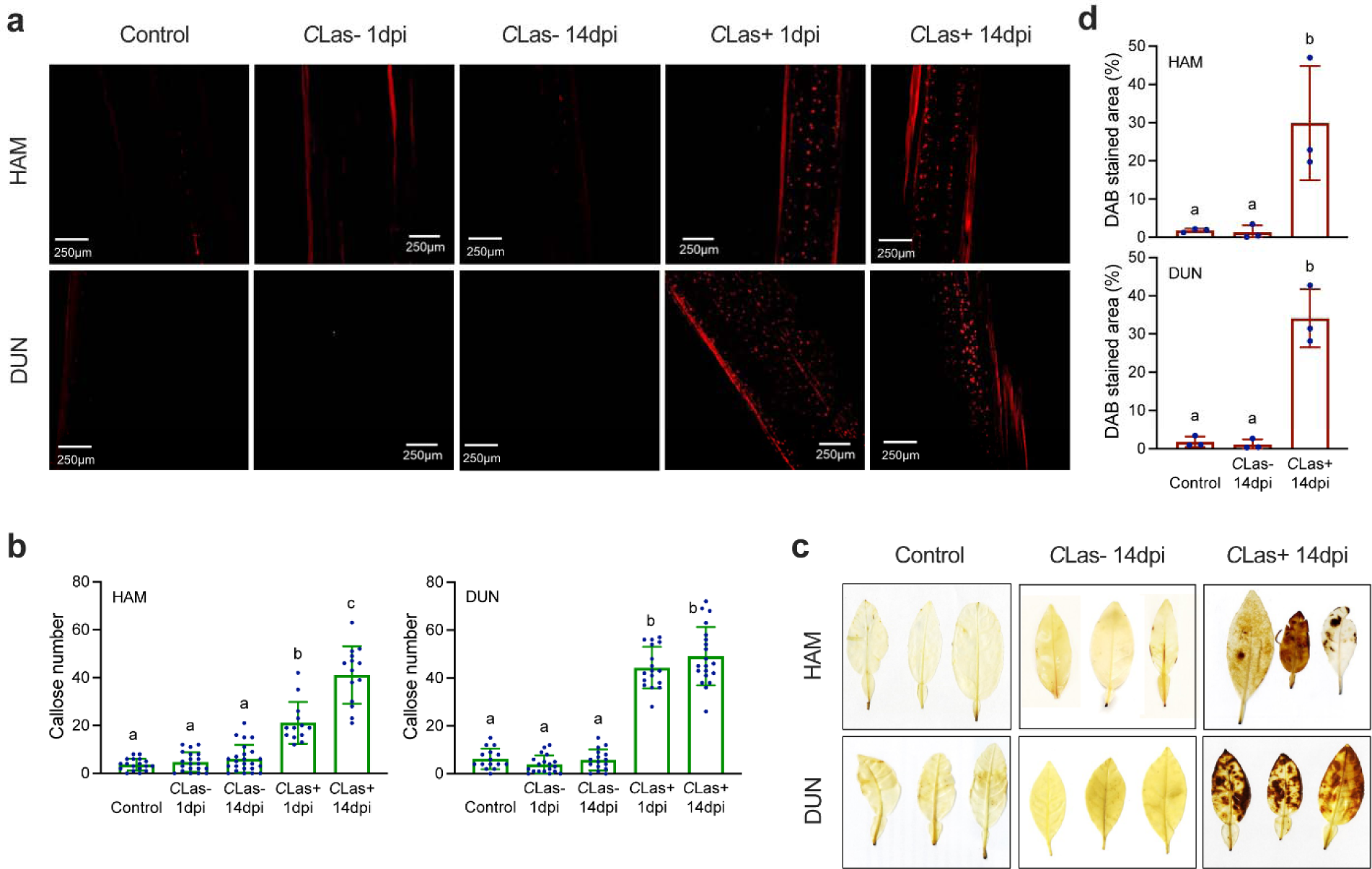
*C*Las-induced callose deposition and ROS accumulation. **a,** Callose deposition (red dots) revealed by aniline blue staining in citrus stems adjacent to the leaves inoculated with *C*Las-free (*C*Las-) or *C*Las-infected (*C*Las+) psyllids at 1 day post-inoculation (dpi) and 14 dpi. Representative images are shown. Control: uninoculated healthy stems; HAM: Hamlin; DUN: Duncan. **b,** Numbers of callose depositions in the control and the citrus stems adjacent to the leaves inoculated with *C*Las-free or *C*Las-infected psyllids at 1 dpi and 14 dpi. Bars represent ‘means ± standard deviation (SD)’ (n = 13-24). Data from three independent experiments were combined. Different letters above the bars denote significant differences (p < 0.05; one-way ANOVA with Tukey’s test). **c,** ROS accumulation (brown precipitates) revealed by DAB staining in citrus leaves inoculated with *C*Las-free or *C*Las-infected psyllids at 14 dpi. Representative images are shown. Control: uninoculated healthy citrus leaves. **d,** Percentages of leaf areas stained with DAB in the control and the citrus leaves inoculated with *C*Las-free or *C*Las-infected psyllids at 14 dpi. Bars represent ‘means ± SD’ (n = 3). Different letters above the bars denote significant differences (p < 0.05; one-way ANOVA with Tukey’s test).

### Overexpression of *AtNPR1* in citrus increases basal callose levels and suppresses *C*Las-induced callose deposition and ROS accumulation

HLB symptom development is almost completely suppressed in the *AtNPR1-OE* citrus plants ^34^. To understand the underlying mechanisms, we inoculated *AtNPR1-OE* and non-transgenic (hereafter wild-type/WT) ‘Hamlin’ and ‘Duncan’ plants with either *C*Las-free or *C*Las-infected psyllids and examined callose and ROS accumulation at 14 dpi. As shown in Fig. 2, *C*Las-free psyllids did not induce callose deposition and ROS accumulation, whereas *C*Las-infected psyllids induced heavy deposition of callose and significant accumulation of ROS in the wild-type citrus plants. Surprisingly, the *AtNPR1-OE* citrus plants exhibited increased basal callose levels in the phloem tissues when uninoculated, and inoculation with either *C*Las-free or *C*Las-infected psyllids did not significantly increase the callose levels (Fig. 2a, b). The callose levels in the *AtNPR1-OE* citrus plants are significantly higher than those in the wild-type when inoculated with *C*Las-free psyllids but are significantly lower than those in the wild-type when inoculated with *C*Las-infected psyllids (Fig. 2a, b). Moreover, the *AtNPR1-OE* citrus plants showed slightly but not significantly increased ROS levels after inoculation with *C*Las-infected psyllids compared to inoculation with *C*Las-free psyllids or without inoculation (Fig. 2c, d). The ROS levels in the *AtNPR1-OE* citrus plants were much lower than those in the wild-type citrus plants after inoculation with *C*Las-infected psyllids (Fig. 2c, d).

**Fig. 2.**
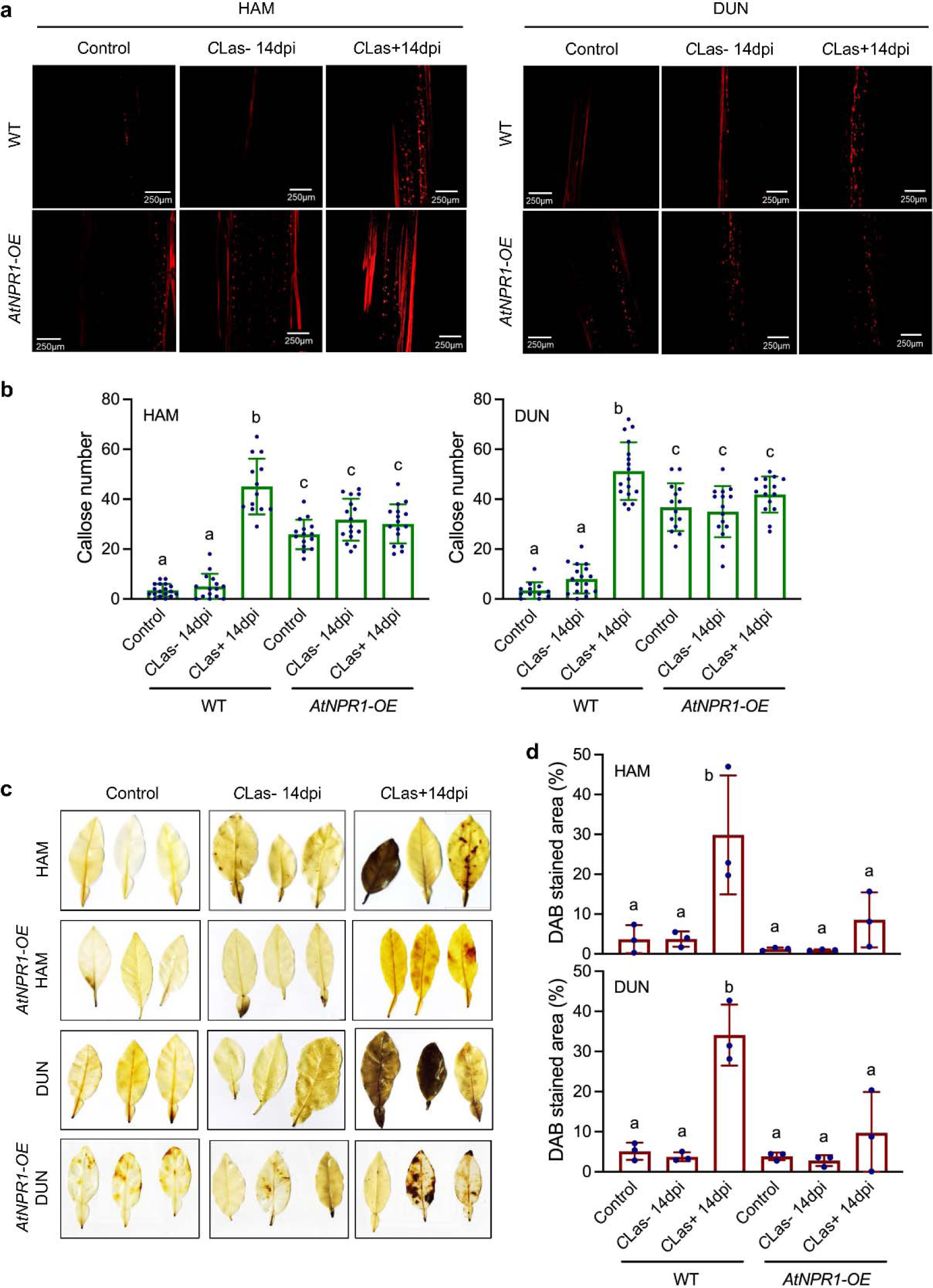
Callose depostion and ROS accumuation in citrus *AtNPR1-OE* plants. **a,** Callose deposition (red dots) revealed by aniline blue staining in wild-type (WT) and *AtNPR1-OE* citrus stems adjacent to the leaves inoculated with *C*Las-free (*C*Las-) or *C*Las-infected (*C*Las+) psyllids at 14 dpi. **b,** Numbers of callose depositions in the wild-type and *AtNPR1-OE* citrus stems adjacent to the leaves inoculated with *C*Las-free or *C*Las-infected psyllids at 14 dpi. Bars represent ‘means ± standard deviation (SD)’ (n = 12-18). Data from three independent experiments were combined. Different letters above the bars denote significant differences (p < 0.05; one-way ANOVA with Tukey’s test). **c,** ROS accumulation (brown precipitates) revealed by DAB staining in wild-type and *AtNPR1-OE* citrus leaves inoculated with *C*Las-free or *C*Las-infected psyllids at 14 dpi. **d,** Percentages of leaf areas stained with DAB in the wild-type and *AtNPR1-OE* citrus leaves inoculated with *C*Las-free or *C*Las-infected psyllids at 14 dpi. Bars represent ‘means ± SD’ (n = 3). Different letters above the bars denote significant differences (p < 0.05; one-way ANOVA with Tukey’s test).

To determine the potential mechanisms underlying the elevated basal levels of callose and the reduced induction of callose deposition and ROS accumulation in the *AtNPR1-OE* citrus plants, we examined the expression levels of *C. sinensis callose synthase 3* (*CsCalS3*) and *CsCalS7* as well as *respiratory burst oxidase homolog D* (*CsRBOHD*) in *AtNPR1-OE* and wild-type ‘Duncan’ plants with or without *C*Las infection. As shown in Fig. 3, the basal transcript levels of *CsCals3* and *CsCals7* are significantly higher in the *AtNPR1*-OE ‘Duncan’ plants than in the wild-type. However, at 14 dpi with *C*Las-infected psyllids, *CsCalS3* and *CsCalS7* transcripts were significantly upregulated in the wild-type ‘Duncan’ but were not induced in the *AtNPR1*-OE ‘Duncan’ plants (Fig. 3). On the other hand, the basal expression level of *CsRBOHD* was significantly lower in the *AtNPR1*-OE plants than in the wild-type (Fig. 3).

**Fig. 3.**
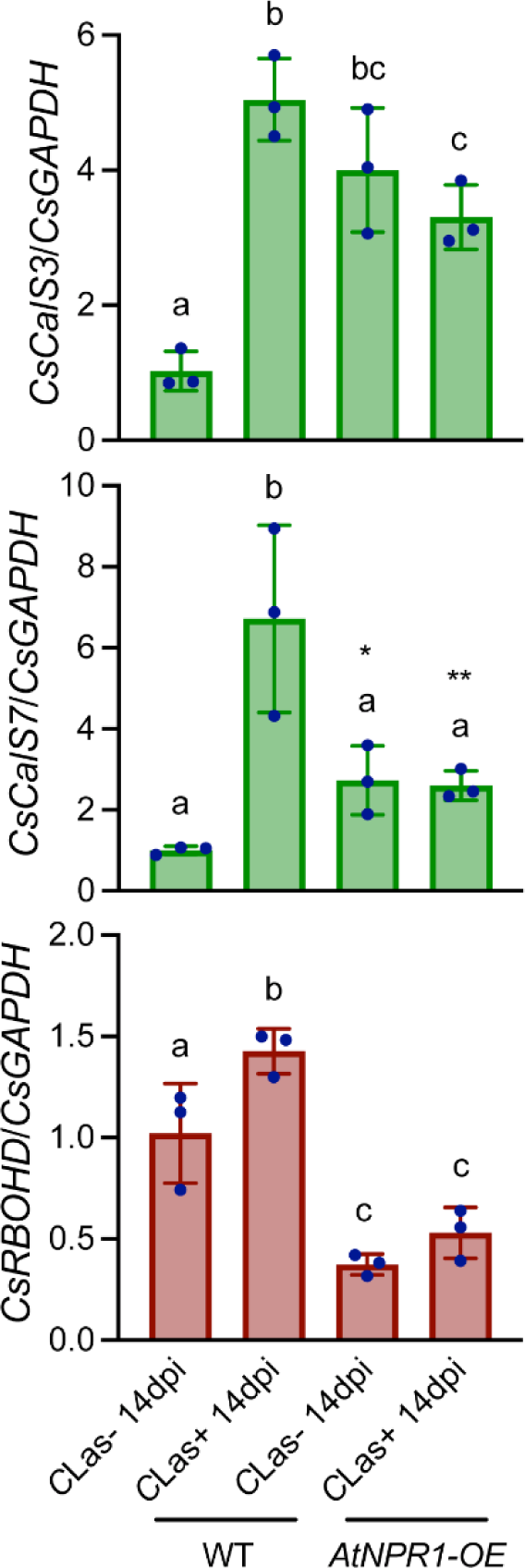
Induction of callose and ROS biosynthesis genes in citrus by *C*Las infection. Expression of *CsCalS3*, *CsCalS7*, and *CsRBOHD* in wild-type (WT) and *AtNPR1-OE* ‘Duncan’ grapefruit leaves inoculated with *C*Las-free (*C*Las-) or *C*Las-infected (*C*Las+) psyllids at 14 dpi. Bars represent ‘means ± standard deviation (SD)’ (n = 3). Different letters above the bars denote significant differences (p < 0.05; one-way ANOVA with Tukey’s test). Asterisks indicate having significant difference from the wild type inoculated with *C*Las-free psyllids (*p < 0.05; **p <0.01: Student’s t-test).

Inoculation with *C*Las-infected psyllids significantly upregulated *CsRBOHD* transcription in the wild-type but had no effect on *CsRBOHD* expression in the *AtNPR1*-OE ‘Duncan’ plants (Fig. 3). These results suggest that *AtNPR1* may modulate callose deposition and ROS accumulation in citrus by regulating the expression of *CsCalS3/7* and *CsRBOHD*, respectively.

### Overexpression of *AtNPR1* in citrus minimizes *C*Las-induced vascular tissue alterations and prevents sieve pore plugging

*C*Las infection causes vascular tissue alterations and sieve pore plugging in susceptible citrus cultivars, resulting in HLB symptom development ^39–41^. To investigate the difference in the vascular bundles between *AtNPR1-OE* and wild-type citrus plants, we carried out thin-section light and transmission electron microscopy (TEM) analysis of the vasculature following inoculation with *C*Las-free or *C*Las-infected psyllids. As shown in Fig. 4a, and 4b, the *AtNPR1-OE* and wild-type ‘Duncan’ plants displayed similar phloem size including new and replacement phloem at 14 dpi with *C*Las-free psyllids. However, the *AtNPR1-OE* ‘Duncan’ plants had larger xylem size than the wild-type (Fig. 4b). Upon inoculation with *C*Las-infected psyllids, we observed a substantial expansion of both phloem and xylem tissues in the wild-type ‘Duncan’ plants, but only a moderate and significantly smaller expansion in the *AtNPR1-OE* ‘Duncan’ plants (Fig. 4a, 4b).

**Fig. 4.**
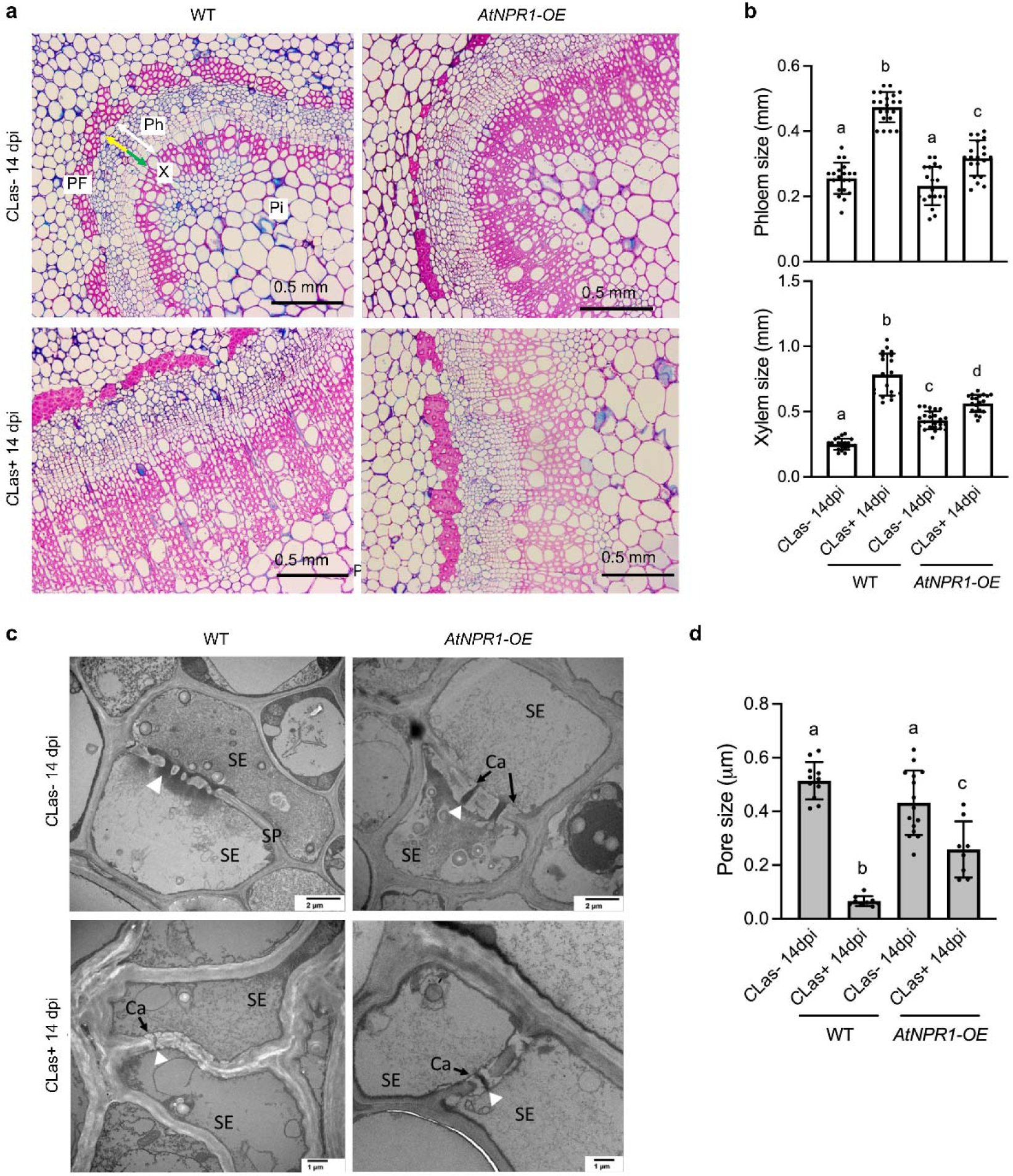
*C*Las-induced vascular tissue alterations and sieve pore plugging in citrus *AtNPR1-OE* plants. **a,** Light microscopy images of methylene blue- and basic fuchsin-stained stem sections of wild-type (WT) and *AtNPR1-OE* ‘Duncan’ grapefruit plants inoculated with *C*Las-free (*C*Las-) or *C*Las-infected (*C*Las+) psyllids at 14 dpi. PF: phloem-fibers; Ph: phloem; X: xylem; Pi: pith; white arrow: phloem size; green arrow: new phloem; yellow arrow: replacement phloem. **b,** The phloem and xylem sizes in the wild-type and *AtNPR1-OE* ‘Duncan’ grapefruit plants inoculated with *C*Las-free or *C*Las-infected psyllids at 14 dpi. Bars represent ‘means ± standard deviation (SD)’ (n = 19-24). Different letters above the bars denote significant differences (p < 0.05; one-way ANOVA with Tukey’s test). **c,** Transmission electron microscopy images of the wild-type and *AtNPR1-OE* ‘Duncan’ grapefruit plants inoculated with *C*Las-free or *C*Las-infected psyllids at 14 dpi. Ca: callose; SP: sieve plate; SE: sieve element; white arrowhead: open passage between two sieve elements. **d,** Pore sizes between sieve elements in the wild-type and *AtNPR1-OE* ‘Duncan’ grapefruit plants inoculated with *C*Las-free or *C*Las-infected psyllids at 14 dpi. Bars represent ‘means ± SD’ (n = 8-15). Different letters above the bars denote significant differences (p < 0.05; one-way ANOVA with Tukey’s test).

The sieve elements of plant stem sections were also observed with TEM. At 14 dpi with *C*Las-free psyllids, we detected a higher amount of callose deposition with visible sieve pore openings in the *AtNPR1-OE* ‘Duncan’ stem sections compared to the wild-type sections that had no significant callose deposition (Fig. 4c, 4d). Upon inoculation with *C*Las-infected psyllids, we observed tremendous callose accumulation in the wild-type ‘Duncan’ stem sections with almost no visible pore openings, whereas the *AtNPR1-OE* ‘Duncan’ plants had much smaller reduction in pore openings caused by callose accumulation (Fig. 4c, 4d).

### Overexpression of *AtNPR1* in *Arabidopsis* increases basal callose levels and suppresses *Psm*-induced callose deposition and ROS accumulation

NPR1 is known to positively contribute to pathogen-induced callose deposition and to negatively regulate pathogen-triggered ROS accumulation ^42–44^. To determine how overexpression of *AtNPR1* influences pathogen-induced callose deposition and ROS accumulation in *Arabidopsis*, we compared the bacterial pathogen *Psm*-induced callose and ROS accumulation in wild-type *Arabidopsis*, *AtNPR1-OE Arabidopsis*, and *npr1-3* mutant plants. Interestingly, the *AtNPR1-OE Arabidopsis* plants accumulated significantly higher basal levels of callose than the wild-type (WT) and *npr1-3*, and *Psm* infection dramatically elevated callose deposition in the wild type but did not significantly change callose levels in the *AtNPR1-OE Arabidopsis* and *npr1-3* plants (Fig. 5a, b). These results revealed a paradox in which the physiologic low level of AtNPR1 is required for *Psm*-induced callose deposition, whereas ectopic expression of high levels of AtNPR1 increases basal callose levels but suppresses *Psm*-induced callose deposition.

**Fig. 5.**
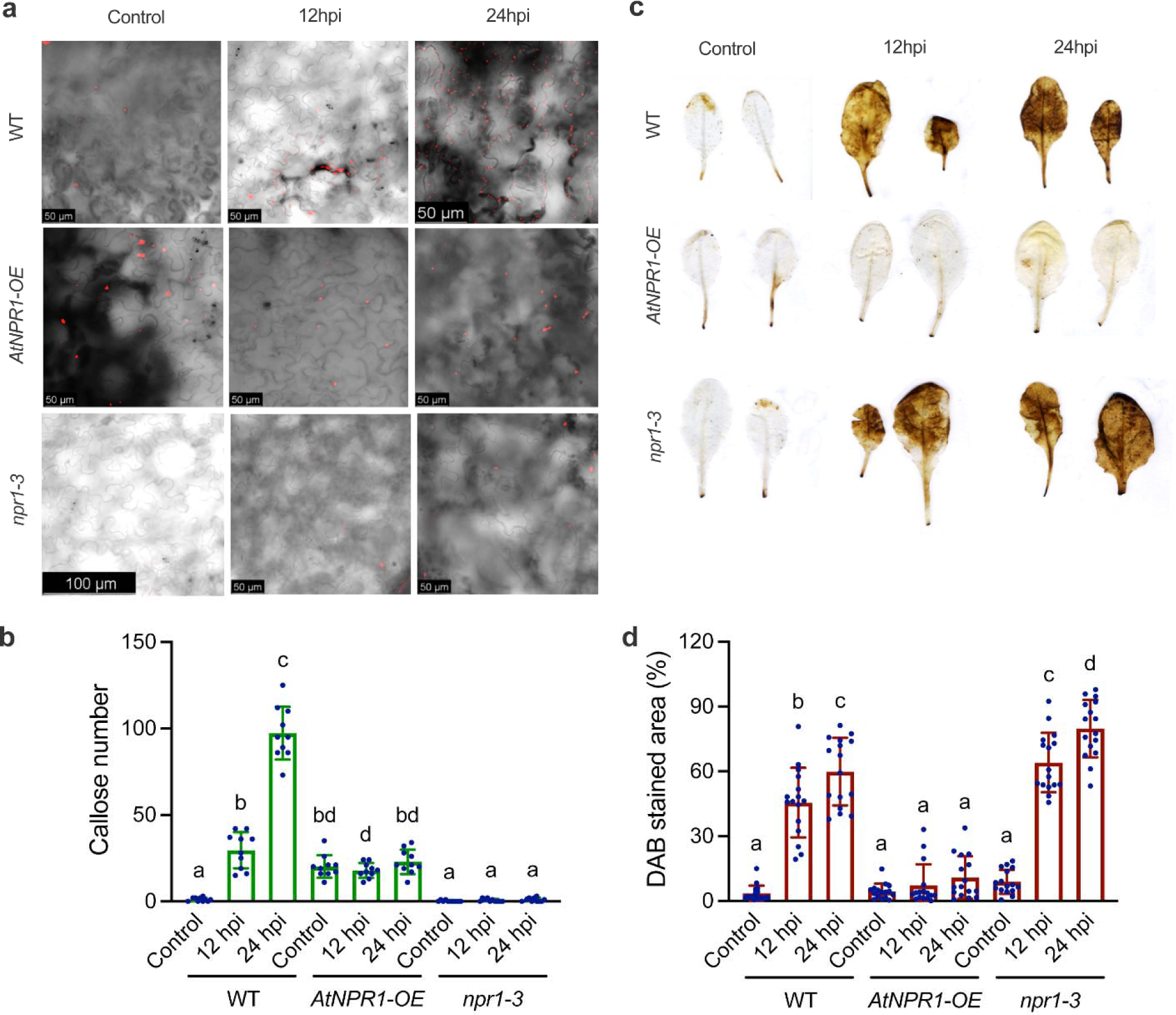
*Psm*-induced callose and ROS accumulation in *Arabidopsis AtNPR1-OE* and *npr1-3* plants. **a,** Callose deposition (red dots) revealed by aniline blue staining in *Arabidopsis* wild-type (WT), *AtNPR1-OE*, and *npr1-3* leaves infected with *Psm* at 12 hr post-inoculation (hpi) and 24 hpi. Representative images are shown. Control: healthy uninfected *Arabidopsis* leaves. **b,** Numbers of callose depositions in the control and the *Arabidopsis* wild-type, *AtNPR1-OE*, and *npr1-3* leaves infected with *Psm* at 12 and 24 hpi. Bars represent ‘means ± standard deviation (SD)’ (n = 10). Data from three independent experiments were combined. Different letters above the bars denote significant differences (p < 0.05; one-way ANOVA with Tukey’s test). **c,** ROS accumulation (brown precipitates) revealed by DAB staining in *Arabidopsis* wild-type, *AtNPR1-OE*, and *npr1-3* leaves infected with *Psm* at 12 and 24 hpi. Representative images are shown. Control: uninoculated healthy *Arabidopsis* leaves. **d,** Percentages of leaf areas stained with DAB in the control and the *Arabidopsis* wild-type, *AtNPR1-OE*, and *npr1-3* leaves infected with *Psm* at 12 and 24 hpi. Bars represent ‘means ± SD’ (n = 16-17). Data from three independent experiments were combined. Different letters above the bars denote significant differences (p < 0.05; one-way ANOVA with Tukey’s test).

Furthermore, *Psm* infection triggered massive ROS accumulation in both the wild-type and *npr1-3* and the ROS levels at 12 hpi are significantly higher in *npr1-3* than those in the wild-type, indicating that AtNPR1 negatively regulates *Psm*-induced ROS accumulation (Fig. 5c, d). In line with this conclusion, *Psm*-induced ROS accumulation was almost completely suppressed in the *AtNPR1-OE Arabidopsis* plants (Fig. 5c, d). These results indicate that AtNPR1 is a major regulator of *Psm*-induced callose deposition and ROS accumulation in *Arabidopsis*.

### Silencing of *CsNPR3* suppresses *C*Las-induced callose and ROS accumulation and confers HLB tolerance

NPR3 and NPR4 are known to negatively regulate plant immunity through NPR1-dependent and NPR1-independent mechanisms. We therefore tested *Psm*-induced callose deposition and ROS accumulation in two *Arabidopsis npr3 npr4* double mutants. Although *Psm* significantly induced callose deposition in the *npr3-1 npr4-3* mutant, the induction level was significantly lower than that in the wild-type plants (Fig. 6a, b), indicating that AtNPR3 and AtNPR4 positively regulate *Psm*-induced callose deposition. Furthermore, both *npr3-2 npr4-2* and *npr3-1 npr4-3* accumulated elevated basal ROS levels and *Psm* infection did not further increase ROS levels in the *npr3-1 npr4-3* mutant (Fig. 6c, d), suggesting that AtNPR3 and AtNPR4 suppress basal ROS and may positively contribute to pathogen-induced ROS accumulation. Since CsNPR3 is the closest citrus ortholog of the AtNPR3 and AtNPR4 proteins (Supplementary Fig. S2), we tested whether the *CsNPR3* gene plays a role in *C*Las-induced callose and ROS accumulation and HLB symptom development. To this end, we used CTV-RNAi to silence the *CsNPR3* gene in citrus. Inoculation of a CTV-tCsNPR3 (truncated *CsNPR3*) construct into *C. macrophylla* efficiently silenced the *CsNPR3* gene (Supplementary Fig. S3). The CTV-tCsNPR3 construct was subsequently graft-inoculated into susceptible ‘Madam Vinous’ sweet orange. ‘Madam Vinous’ plants carrying the wild-type CTV (CTV-wt) and the CTV-tCsNPR3 construct were inoculated with *C*Las-infected psyllids. After becoming *C*Las positive, these plants were kept in greenhouse for symptom development. Nine months later, the ‘Madam Vinous’ plants carrying CTV-wt all exhibited severe HLB symptoms, whereas those with CTV-tCsNPR3 displayed segregating phenotype with no or mild symptoms (Fig. 7a, b). We then determined callose and ROS levels in these plants and the control plants (with or without CTV and no *C*Las). CTV-wt significantly and moderately increased ROS levels in the plants (Fig. 8a-d). Interestingly, silencing of *CsNPR3* elevated basal callose levels and suppressed *C*Las-induced callose deposition and ROS accumulation (Fig. 8a-d). We next propagated the *CsNPR3* RNAi lines by grafting to test whether the HLB tolerance could be retained in the progeny. Nine months after the propagation, all progeny plants showed no or mild symptoms (Fig. 7c, d). These results together indicate that silencing of *CsNPR3* created HLB tolerance in susceptible ‘Madam Vinous’ by repressing *C*Las-induced overaccumulation of callose and ROS. Thus, *CsNPR3* plays a positive role in HLB disease symptom development.

**Fig. 6.**
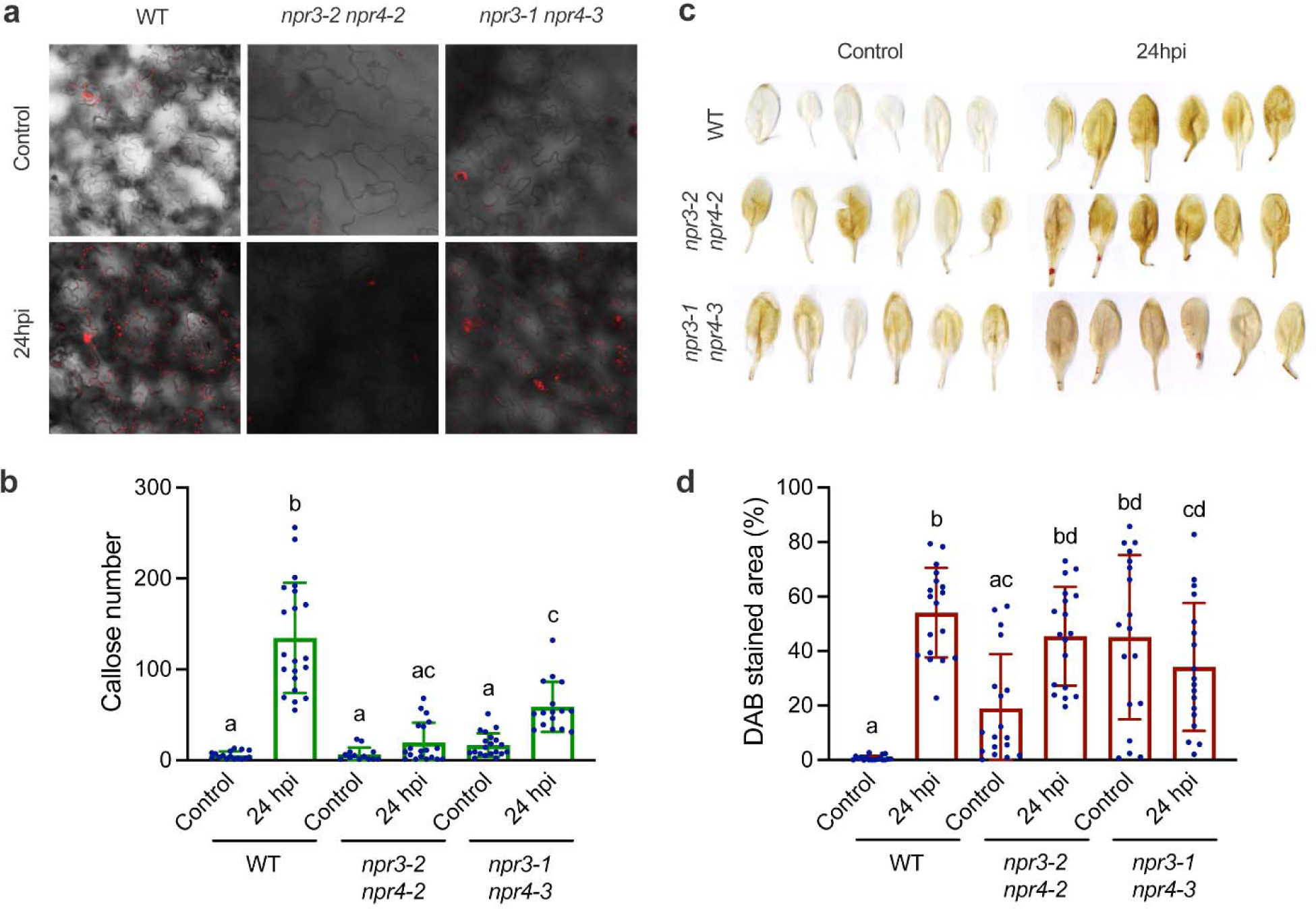
*Psm*-induced callose and ROS accumulation in *Arabidopsis npr3 npr4* plants. **a,** Callose deposition (red dots) revealed by aniline blue staining in *Arabidopsis* wild-type (WT), *npr3-2 npr4-2*, and *npr3-1 npr4-3* leaves infected with *Psm* at 24 hpi. Representative images are shown. Control: healthy uninfected *Arabidopsis* leaves. **b,** Numbers of callose depositions in the control and the *Arabidopsis* wild-type, *npr3-2 npr4-2*, and *npr3-1 npr4-3* leaves infected with *Psm* at 24 hpi. Bars represent ‘means ± standard deviation (SD)’ (n = 13-21). Data from three independent experiments were combined. Different letters above the bars denote significant differences (p < 0.05; one-way ANOVA with Tukey’s test). **c,** ROS accumulation (brown precipitates) revealed by DAB staining in *Arabidopsis* wild-type, *npr3-2 npr4-2*, and *npr3-1 npr4-3* leaves infected with *Psm* at 24 hpi. Representative images are shown. Control: uninoculated *Arabidopsis* leaves. (d) Percentages of leaf areas stained with DAB in the control and the *Arabidopsis* wild-type, *npr3-2 npr4-2*, and *npr3-1 npr4-3* leaves infected with *Psm* at 24 hpi. Bars represent ‘means ± SD’ (n = 18). Data from three independent experiments were combined. Different letters above the bars denote significant differences (p < 0.05; one-way ANOVA with Tukey’s test).

**Fig. 7.**
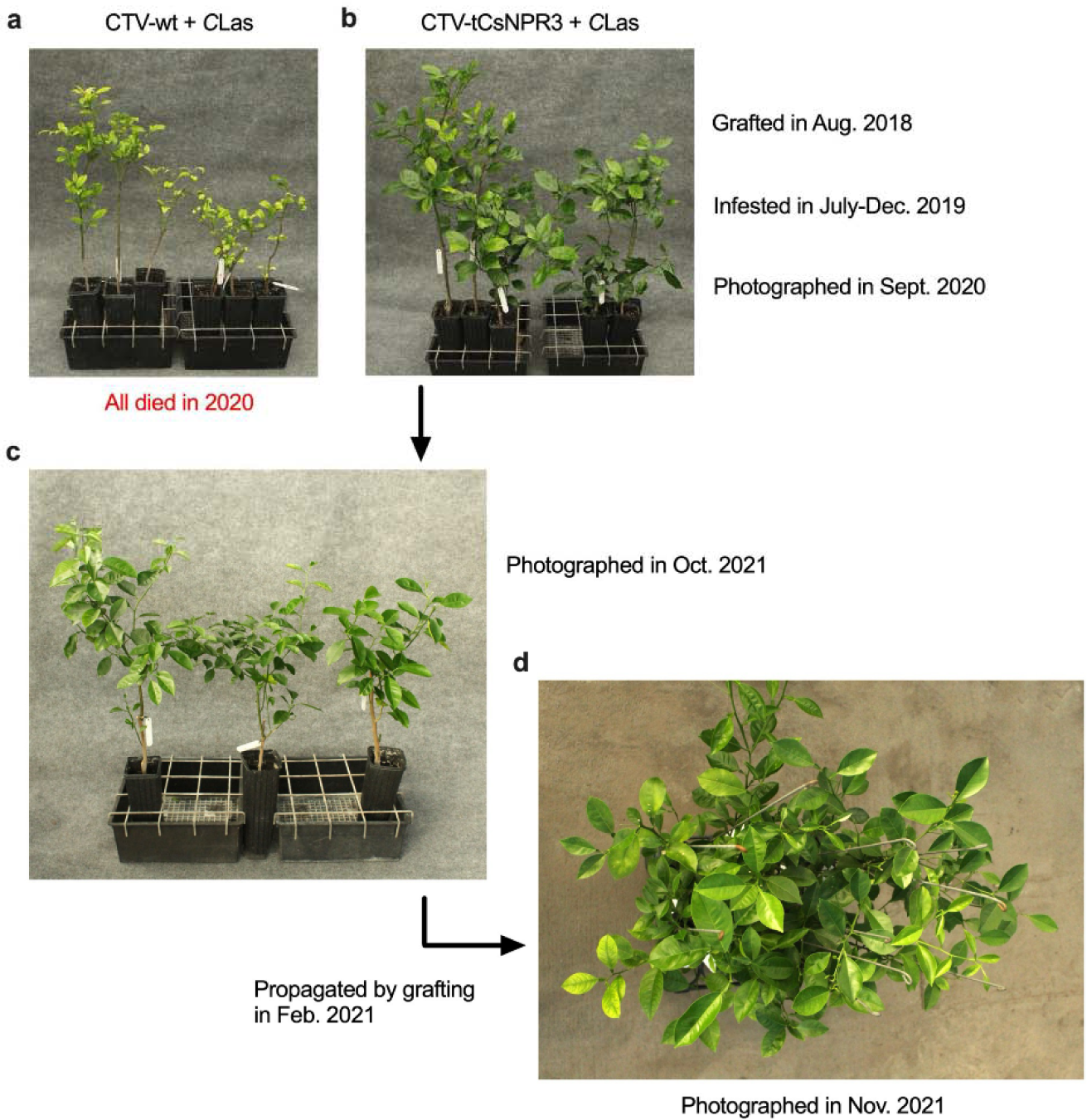
HLB symptom development in *C*Las-infected citrus *CsNPR3* RNAi plants. The wild-type CTV (CTV-wt) and CTV-tCsNPR3 (truncated *CsNPR3*) constructs were graft-inoculated into susceptible ‘Madam Vinous’ sweet orange in August 2018. The ‘Madam Vinous’ plants carrying the CTV constructs were infested with *C*Las-infected psyllids from July to December 2019. Photos were taken in September 2020. All CTV-wt plants infected by *C*Las died in 2020. The *C*Las-positive CTV-tCsNPR3 plants (*CsNPR3* RNAi lines) were propagated in February 2021. Photos of the parents and progenies were taken in November 2021. **a,** HLB symptoms on *C*Las-positive ‘Madam Vinous’ plants carrying CTV-wt. **b,** HLB symptoms on *C*Las-positive ‘Madam Vinous’ plants carrying CTV-tCsNPR3. **c,** The parental *C*Las-positive CTV-tCsNPR3 plants that provided budwoods for propagation. **d,** HLB symptoms on the *C*Las-positive CTV-tCsNPR3 progeny plants.

**Fig. 8.**
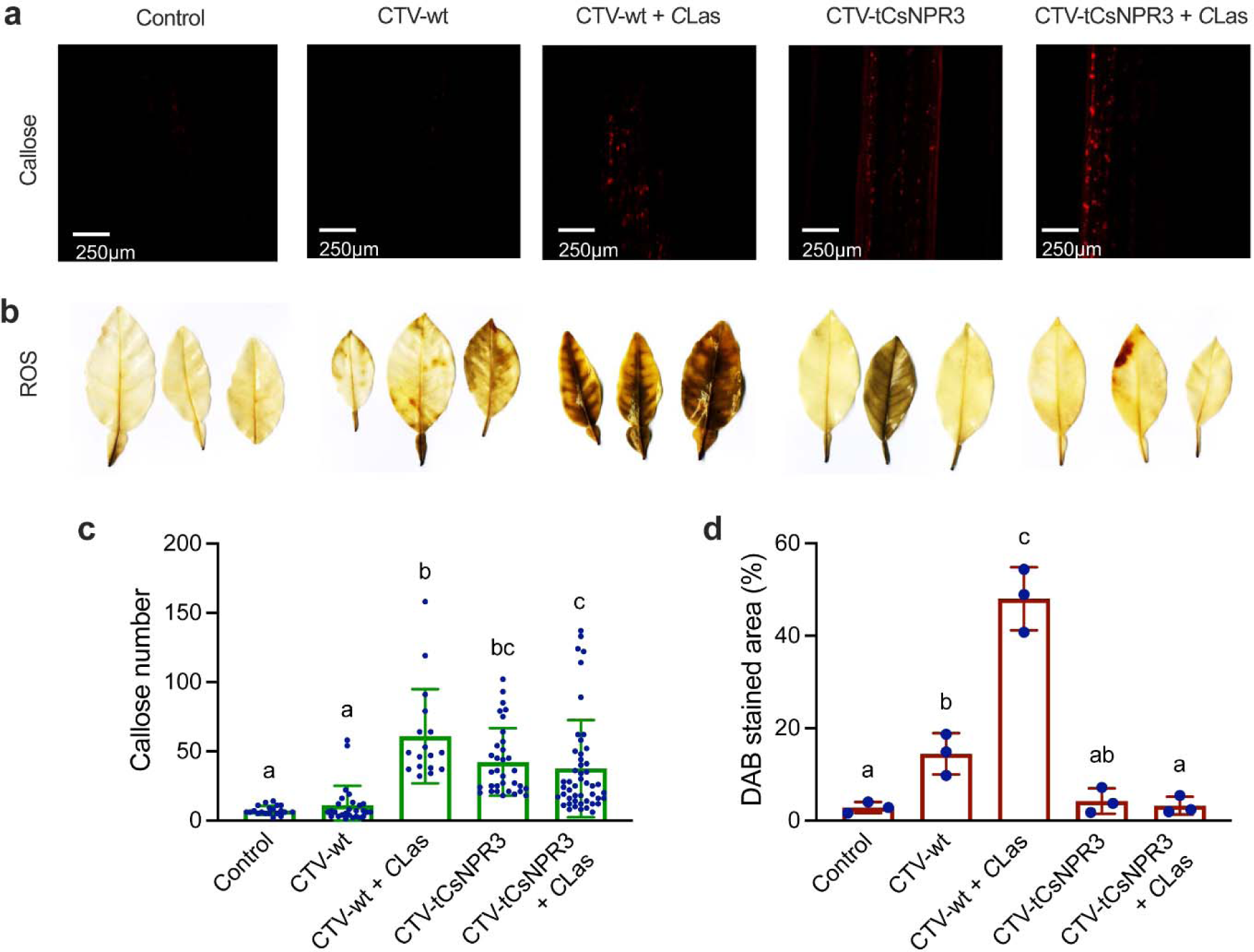
*C*Las-induced callose and ROS accumulation in citrus *CsNPR3* RNAi plants. **a,** Callose deposition (red dots) revealed by aniline blue staining in CTV-wt, *C*Las-positive CTV-wt, CTV-tCsNPR3, and *C*Las-positive CTV-tCsNPR3 ‘Madam Vinous’ stems. Representative images are shown. Control: healthy ‘Madam Vinous’ stems. **b,** ROS accumulation (brown precipitates) revealed by DAB staining in CTV-wt, *C*Las-positive CTV-wt, CTV-tCsNPR3, and *C*Las-positive CTV-tCsNPR3 ‘Madam Vinous’ leaves. Control: healthy ‘Madam Vinous’ leaves. **c,** Numbers of callose depositions in the control and the CTV-wt, *C*Las-positive CTV-wt, CTV-tCsNPR3, and *C*Las-positive CTV-tCsNPR3 ‘Madam Vinous’ stems. Bars represent ‘means ± standard deviation (SD)’ (n = 17-48). Data from three independent experiments were combined. Different letters above the bars denote significant differences (p < 0.05; one-way ANOVA with Tukey’s test). **d,** Percentages of leaf areas stained with DAB in the control and the CTV-wt, *C*Las-positive CTV-wt, CTV-tCsNPR3, and *C*Las-positive CTV-tCsNPR3 ‘Madam Vinous’ leaves. Bars represent ‘means ± SD’ (n = 3). Different letters above the bars denote significant differences (p < 0.05; one-way ANOVA with Tukey’s test).

## Discussion

*C*Las infection of most commercial citrus varieties triggers chronic immune imbalance resulting in overaccumulation of callose and ROS in the phloem ^4,45^. High levels of callose and ROS have been shown to cause phloem plugging ^4^, which hinder the export of photoassimilates, resulting in accumulation of starch and development of HLB symptoms ^4,6,39,41^. Interestingly, some citrus varieties and relatives, e.g., ‘Carrizo’ citrange and ‘Persian’ lime, exhibit little or no HLB symptoms after being infected by *C*Las, a phenomenon termed HLB tolerance ^46,47^. Tolerant citrus varieties display increased induction of genes which encode key immune regulators and enzymes that can break down callose and ROS, preventing their overaccumulation ^15^. These results suggest that the HLB-susceptible varieties lack the ability to maintain the immune balance needed to prevent *C*Las-induced overaccumulation of callose and ROS. Although the genetic defect(s) in the immune systems of susceptible varieties is currently unclear, transgenic studies have shown that overexpression of the *AtNPR1* gene confers robust HLB tolerance in susceptible varieties including ‘Hamlin’ sweet orange and ‘Duncan’ grapefruit ^34^. We further show here that overexpression of *AtNPR1* elevates basal callose levels and suppresses *C*Las-induced callose and ROS accumulation (Fig. 2). Importantly, we found that silencing of *CsNPR3*, which encodes a negative regulator of the SA and NPR1 signaling pathway, in susceptible ‘Madam Vinous’ sweet orange similarly increases basal callose levels, inhibits *C*Las-induced callose and ROS accumulation, and enhances HLB tolerance (Fig. 7 and Fig. 8). These results indicate that the immune balance of susceptible varieties in response to *C*Las infection can be restored by overexpression of *AtNPR1* or silencing of *CsNPR3* and suggest that the immune defect(s) in HLB-susceptible varieties can be permanently cured by tipping the immune balance via genetic approaches.

ROS plays an important signaling role in plant immunity; however, high levels of ROS often cause oxidative stress damaging cellular components ^48^. ROS accumulation is therefore tightly regulated through the interplay between ROS production and scavenging. The plasma membrane-localized RBOHs are the major enzymes mediating pathogen-induced ROS production ^49^. The activity of RBOHs is precisely controlled by positive and negative regulatory mechanisms that are concomitantly activated in response to pathogen infection. For instance, phosphorylation of the *A. thaliana* RBOHD (AtRBOHD) at Ser 343, Ser347, and Ser703 increases its activity, whereas phosphorylation at Thr912 enhances its degradation ^50–52^. Moreover, production of *S*-nitrosothiols (SNOs) by addition of nitric oxide moieties to cysteine thiols facilitates ROS production; however, high levels of SNOs triggers *S*-nitrosylation of AtRBOHD at Cys890, which inhibits its ROS-synthesizing activity ^53^. ROS can also be removed by antioxidant enzymes including catalase, glutathione/ascorbate peroxidase, and glutathione S-transferase ^54^. The activities of these enzymes are significantly upregulated upon pathogen infection, which ensures that ROS subside after the initial oxidative bursts ^55^. Thus, *C*Las-triggered overaccumulation of ROS in susceptible citrus varieties is likely a result of impaired negative regulation of ROS production and/or compromised induction of antioxidant enzymes.

ROS are known to positively contribute to elicitor-induced callose deposition ^1^. For instance, callose deposition induced by several pathogen-associated molecular patterns is compromised in the ROS-defective mutants *peroxidase33* (*prx33*) *prx34* and *rbohD* ^56^. The *rbohD* mutant also exhibits reduced callose deposition elicited by oligogalacturonides ^57^. It has been shown that *C*Las induces ROS accumulation earlier than callose deposition in new shoots on *C*Las-positive plants ^4^, suggesting that ROS might facilitate callose deposition in HLB symptom development. However, when *C*Las is transmitted into new shoots on naïve healthy plants by psyllids, callose deposition is induced long before ROS accumulation (Fig. 1), suggesting that *C*Las-induced initial callose deposition is largely independent of ROS accumulation. Interestingly, *C*Las can reduce callose levels at the phloem sieve tubes to facilitate its intercellular movement ^58^, suggesting that *C*Las-induced callose deposition is an effective defense mechanism against this bacteria in citrus. However, overaccumulation of callose can cause phloem plugging, contributing to HLB symptom development. It is well known that callose levels are determined by the equilibrium between synthesis and removal ^59^. For instance, SA treatment and pathogen infection in *Arabidopsis* simultaneously induce the expression of callose biosynthetic genes including *AtCalS1* and *AtCalS12* as well as catabolic genes such as *PR2* that encodes a β-1,3-glucanase degrading callose ^60^. Furthermore, callose deposition in the phloem, especially in the sieve elements, is greatly reduced in *cals7* mutants ^61^, and enhancement of *AtCalS3* expression during phloem development is able to complement the *cals7* defects ^62^. Thus, *C*Las-induced overaccumulation of callose in the phloem may result from heightened induction of callose biosynthetic genes and/or compromised induction of catabolic genes.

NPR1 has been shown to play a significant role in regulating ROS production and scavenging. For instance, a strong oxidative burst occurs in *npr1* roots but not in wild-type roots upon inoculation with *Sinorhizobium meliloti* or treatment with a purified nodulation factor ^63^. Similarly, higher levels of ROS and lipid peroxidation were detected in *npr1* systemic tissues than in wild-type during SAR induction ^64^. Moreover, hydroxyl radicals cause more severe damage to *npr1* seedling roots than to wild-type ^65^, and *npr1* seedlings are hypersensitive to ROS-mediated SA toxicity ^66–70^. On the other hand, *Arabidopsis* plants overexpressing the mulberry *NPR1* (*MuNPR1*) gene and potato (*Solanum tuberosum*) plants overexpressing *StNPR1* exhibit decreased H_2_O_2_ and/or O_2_^-^ levels in response to *P. syringae* pv. *tomato* (*Pst*) DC3000 and *Rastonia solanacearum* infection, respectively, probably owing to increased ROS scavenging activities ^71,72^. Furthermore, *Nicotiana tabacum* plants overexpressing *AtNPR1* display enhanced tolerance to methyl viologen and accumulate reduced levels of ROS in chloroplasts under salt stress, likely due to upregulated expression of genes encoding ROS-scavenging enzymes ^44,73,74^ The *npr1* mutant accumulates higher levels of ROS than wild-type in response to *Psm*, whereas overexpression of *AtNPR1* in *Arabidopsis* and citrus drastically inhibits *Psm*- and *C*Las-induced ROS accumulation, respectively (Fig. 2c, 2d and Fig. 5c, 5d). Together these results demonstrate that NPR1 can prevent pathogen-induced overaccumulation of ROS and alleviate the toxic effects of excessive ROS.

NPR1 appears to play a dual role in pathogen-induced callose deposition ^1,42,43,75^. In *Arabidopsis*, *AtCalS1* and *AtCalS12* are highly induced by SA treatment, the presence of *Hyaloperonospora arabidopsis*, or *Pst* DC3000 Δ*avrPtoB*, and the induction is significantly reduced in the *npr1* mutant plants ^42,43,75^ (Fig. 5). These results indicate that NPR1 is a positive regulator of callose deposition. Consistent with this conclusion, *AtNPR1-OE Arabidopsis* and citrus plants accumulate elevated basal callose levels (Fig. 2a, 2b and Fig. 5a, 5b). Interestingly, *Psm*- and *C*Las-induced callose deposition is significantly diminished in the *AtNPR1-OE Arabidopsis* and citrus plants, respectively (Fig. 2a, 2b and Fig. 5a, 5b). Although the molecular mechanism underlying these new observations remains to be uncovered, diminishing *C*Las-induced callose deposition likely contributes to the HLB tolerance of the transgenic plants.

*C*Las infection in wild-type ‘Duncan’ plants caused a dramatic increase in phloem and xylem sizes at 14 dpi, but a much smaller increase was observed in the *AtNPR1-OE* ‘Duncan’ plants (Fig. 4a, b). A similar observation was shown in the vascular bundles of midribs of healthy and *C*Las-infected susceptible ‘Pineapple’ sweet orange and tolerant ‘Sugar Belle’ mandarin, where phloem and xylem sizes increased after infection in the susceptible variety, but not in the tolerant variety ^76^. We hypothesize that the plants are trying to overcome the callose-plugged phloem by producing more vasculature, resulting in enhanced vascular regeneration. In support of this hypothesis, a similar scenario takes place in the stem pitting disease caused by CTV ^77^.

NPR3 and NPR4 are SA receptors, acting as either adaptor proteins of the CUL3 E3 ligase that specifically target NPR1 for degradation or transcriptional corepressors of defense genes such as *SAR DEFICIENT1* and *WRKY70* ^28,29,78^. It has been shown that SA-induced upregulation of *AtRBOHD* and *AtRBOHF* is compromised in the *npr3 npr4* double mutant ^79^ and *Arabidopsis* plants overexpressing the *MuNPR4* gene accumulate increased levels of H_2_O_2_ and O_2_^-^ after *Pst* DC3000 infection ^72^. We found that basal ROS levels are elevated and *Psm*-induced callose deposition and ROS accumulation are inhibited in *npr3 npr4* mutant plants (Fig. 6). These results indicate that NPR3 and NPR4 positively contribute to pathogen-induced callose and ROS accumulation. In line with this conclusion, silencing of *CsNPR3* mimics overexpression of *AtNPR1*, increasing basal callose levels, suppressing *C*Las-induced callose and ROS accumulation, and inhibiting HLB symptom development (Fig. 7 and Fig. 8). Although the specificity of the *CsNPR3* RNAi remains to be determined, these results indicate that *CsNPR3* plays a positive role in HLB symptom development and suggest that HLB tolerance can be achieved by silencing or knocking out negative regulators, e.g., *CsNPR3*, of the SA and NPR1 signaling pathway in citrus.

Overexpression of *AtNPR1* or its orthologs has been shown to enhance resistance to a broad spectrum of pathogens including bacterial, fungal, and viral pathogens in numerous crop species such as rice, wheat, cotton, soybean, potato, tomato, strawberry, peanut, apple, and grape ^80^. These findings demonstrate the value of *NPR1* as a transgenic crop protection strategy against diverse diseases. Although overexpression of *AtNPR1* does not reduce the titer of *C*Las in citrus plants ^34^, it suppresses *C*Las-induced callose deposition and ROS accumulation (Fig. 2), leading to HLB tolerance. Since overexpression of *AtNPR1* or its orthologs in other plant species also represses pathogen-induced callose and/or ROS accumulation ^44,71–74^ (Fig. 5), this previously overlooked function of *NPR1* is conserved ^81^. By repressing pathogen-induced callose and ROS accumulation, overexpression of *NPR1* in citrus and other crops can hinder the disease development and provide a more sustainable and durable protection. Therefore, creating HLB tolerance in susceptible citrus varieties by overexpression of *AtNPR1* is a highly reliable and promising approach to mitigating the HLB disease.

## Methods

### Plant materials

Budwood of the previously reported *AtNPR1*-OE ‘Hamlin’ sweet orange line 13-3 and ‘Duncan’ grapefruit line 57-28 ^32^ and the corresponding non-transgenic wild types were grafted onto ‘Volkamer’ lemon rootstock using a wedge-grafting method. All progeny plants were grown in a greenhouse with controlled temperature and humidity of 25°C and 60%, respectively. The grafted plants were used two months later for *C*Las inoculation experiments. *Arabidopsis* plants used in this study are wild-type Col-0, *AtNPR1-OE* in Col-0 background ^18^, *npr1-3* ^66^, *npr3-2 npr4-2*, and *npr3-1 npr4-3* ^37^.

To create *CsNPR3* RNAi plants, a cDNA fragment of *CsNPR3* was amplified with the primers *StuI-tCsNPR3F* and *PacI-tCsNPR3R* (Supplementary Table 1) and cloned into pCAMBIA1380 with p22 of ToCV using PacI and StuI at the 3’ end of CTV under the duplicated CP-CE of CTV as previously described ^38^. The empty vector was used as a control (CTV-wt). The constructs were introduced into *N. benthamiana* and subsequently bark-snap inoculated into *C. macrophylla* as described ^38^. After systemic infection in the *C. macrophylla* plants was confirmed by enzyme-linked immunosorbent assay (ELISA) and silencing of *CsNPR3* was validated by quantitative PCR (qPCR), the constructs were graft-transmitted into ‘Madam Vinous’ sweet orange. After systemic infection in the ‘Madam Vinous’ plants was verified by ELISA, the plants were inoculated with *C*Las-infected psyllids in a containment growth room as described previously ^34^. Once the plants were tested *C*Las positive, they were maintained in a clean growth room for HLB symptom development.

### Insects

Populations of healthy and *C*Las-infected Asian citrus psyllids (*Diaphorina citri*) were maintained in cages in controlled rooms on curry leaf plants (*Bergera koenigii*) and *C. macrophylla* plants, respectively. The populations were tested periodically for the presence of *C*Las by qPCR with *C*Las-specific primers (Supplementary Table 1).

### *C*Las and *Psm* infection

For *C*Las infection of citrus, a single leaf, still attached to each plant was placed in a closed chamber (a clear plastic cup with a dome lid) and *C*Las-free or *C*Las-infected psyllids were released into the chamber to feed on the leaf. The leaves and stem-phloem bark adjacent to the inoculated leaves were harvested at the indicated time points for callose and DAB staining as well as gene expression. For *Psm* infection of *Arabidopsis*, an overnight culture of the bacterial pathogen *Psm* with an OD_600_ of 0.2 was used. The culture was diluted in 1 mM MgCl_2_ to an OD_600_ of 0.001 and the diluted *Psm* suspension was infiltrated into fully expanded leaves on four-week-old *Arabidopsis* plants. The 1mM MgCl_2_ solution was used as mock inoculation control. The infiltrated leaves were collected at 12 and 24 hpi for callose and ROS detection.

### Callose detection

For callose detection in the phloem, the stem bark adjacent to the psyllid-infested leaves was peeled (1-2 cm long) using a sharp scalpel, ensuring that the phloem portion was included. At least three citrus trees were used as biological replicates. The samples were snap chilled in 85% ice-cold ethanol followed by fixation overnight in the same solution at room temperature. For *Arabidopsis*, the *Psm*-infiltrated leaves were detached from the plants at 12 and 24 hpi and dipped in chilled 85% ethanol followed by overnight fixation. The ethanol was removed the next day, and the samples were washed with 0.01 % Tween 20 for 1 hr, followed by staining with 0.1% aniline blue (in 10 mM potassium phosphate buffer, pH 12) in the dark for at least 1 hr. The samples were visualized under Leica SP8 confocal microscope with an excitation wavelength of 405 nm and a band-pass 420-480 nm emission filter. The number of callose deposits per 10× (40× for *Arabidopsis*) field was detected using a modified macrocode in ImageJ software ^82,83^.

### Detection of ROS

The infected leaves were collected and immediately submerged in a freshly made DAB solution (1 mg/mL, pH 3.0) and incubated overnight on a shaker at 50 rpm. The leaf samples were washed twice in a destaining solution (ethanol/glycerol/acetic acid, ratio 3/1/1), each for 20 min at 95°C and room temperature, respectively. The leaf samples were scanned, and the staining area was measured in ImageJ using the Color Devonvolution2 plugin and the H DAB vector. The percentage of leaf area with brown stain was calculated using the equation: % DAB-stained area = (stained leaf area/total leaf area) × 100.

### RNA extraction and quantitative PCR (qPCR)

Leaf samples (100 mg) were detached and immediately snap frozen in liquid N_2_ and ground in a tissue lyser for RNA isolation using TRI RNA Isolation Reagent (Sigma) according to the manufacturer’s instructions. The isolated RNA was used for cDNA preparation using High-Capacity cDNA Reverse Transcription Kit (Applied Biosystems) according to the manufacturer’s instruction. qPCR was performed using SYBR™ Green PCR Master Mix (Applied Biosystems) on a 7500 Fast Real-Time PCR system (Applied Biosystems) according to the user’s manual. The 2^-ΔΔCt^ method was used to determine the relative level of gene expression ^84^. The *CsGAPDH* gene was used as an internal control. The efficiency of the primers were determined as previously described ^85^. The primers are listed in Supplementary Table 1.

### Light microscopy and transmission electron microscopy

For both light microscopy and electron microscopy, the stems of each sample were cut into 1 cm pieces and immediately fixed in 3% (v/v) glutaraldehyde in 0.1 M potassium phosphate buffer (pH 7.2) for 4 hr at room temperature. After washing with more phosphate buffer, the samples were fixed again in 2% osmium tetraoxide for 4 hr at room temperature. The samples were again washed with the same phosphate buffer. The samples were next dehydrated in a series of acetone solutions, starting at 10% and ending at 100%, increasing by 10% each stage, soaking in each solution for 10 minutes. The samples were placed in a 50% (v/v) Spurr’s resin and acetone solution, then a 70% (v/v) Spurr’s resin and acetone solution, each for 8 hours. Lastly, the samples were embedded in 100% Spurr’s resin by baking in a 70°C oven overnight. For light microscopy, the samples were semithin-sectioned at a depth of 0.99 µm. The sections were placed on a glass slide and heat sealed on a slide warmer. The cells in the section were stained with methylene blue followed by counter stain in basic fuchsin to stain lignins and other hydrophobic compounds in the cells for 30 sec each. The dried and mounted sections were imaged with a light microscope with an attached camera. The images were analyzed in ImageJ for measurement of phloem and xylem sizes. For electron microscopy, ultrathin sections of 90 nm were made and mounted onto 200-mesh formvar-coated copper grids, stained with Uranyless for 10 minutes, rinsed with distilled water, and stained with Reynold’s lead citrate for 5 minutes. The grids were washed with distilled water and 0.1 N NaOH, then allowed to dry overnight. The samples were then observed with a Hitachi H-7650 transmission electron microscope with an attached AMT XR50 camera. Images obtained from the observations were analyzed in ImageJ and the sieve plate openings were measured.

### Statistical analysis

Each experiment was repeated at least three times. Statistical analyses were performed using the data analysis tools (Student’s t-test) in Microsoft Excel of Microsoft Office 2023 for Macintosh as well as one-way ANOVA in Prism 10.

## Data availability

The authors declare that all data supporting the findings of this study are available within the manuscript and its supplementary files or are available from the corresponding authors upon request. Source data are provided with this paper.

## Acknowledgements

The authors would like to thank Chunxia Wang for comments and suggestions, Dr. Qi Li for growing *Arabidopsis* plants, and Turksen Shilts for conducting the qPCR on *Cs*NPR3-RNAi plants. This research was supported by the National Institute of Food and Agriculture (Grant 2020-70029-33197) to A.L. and Z.M.

## Author contributions

P.S., A.L., and Z.M. conceived the study. P.S. conducted pathogen infection, callose and DAB staining, and gene expression experiments, as well as data analysis. S.W. performed the TEM experiment. C.E.-M. and C.J.R. carried out the CTV RNAi experiments. D.T. propagated the citrus plants. V.O. generated the transgenic plants. P.S., A.L., and Z.M. wrote the paper with input from all authors.

