## Supplementary material for "*NONEXPRESSOR OF PATHOGENESIS-RELATED GENES* control Huanglongbing tolerance by regulating immune balance in citrus plants": Fig. S

**Supplementary Information**

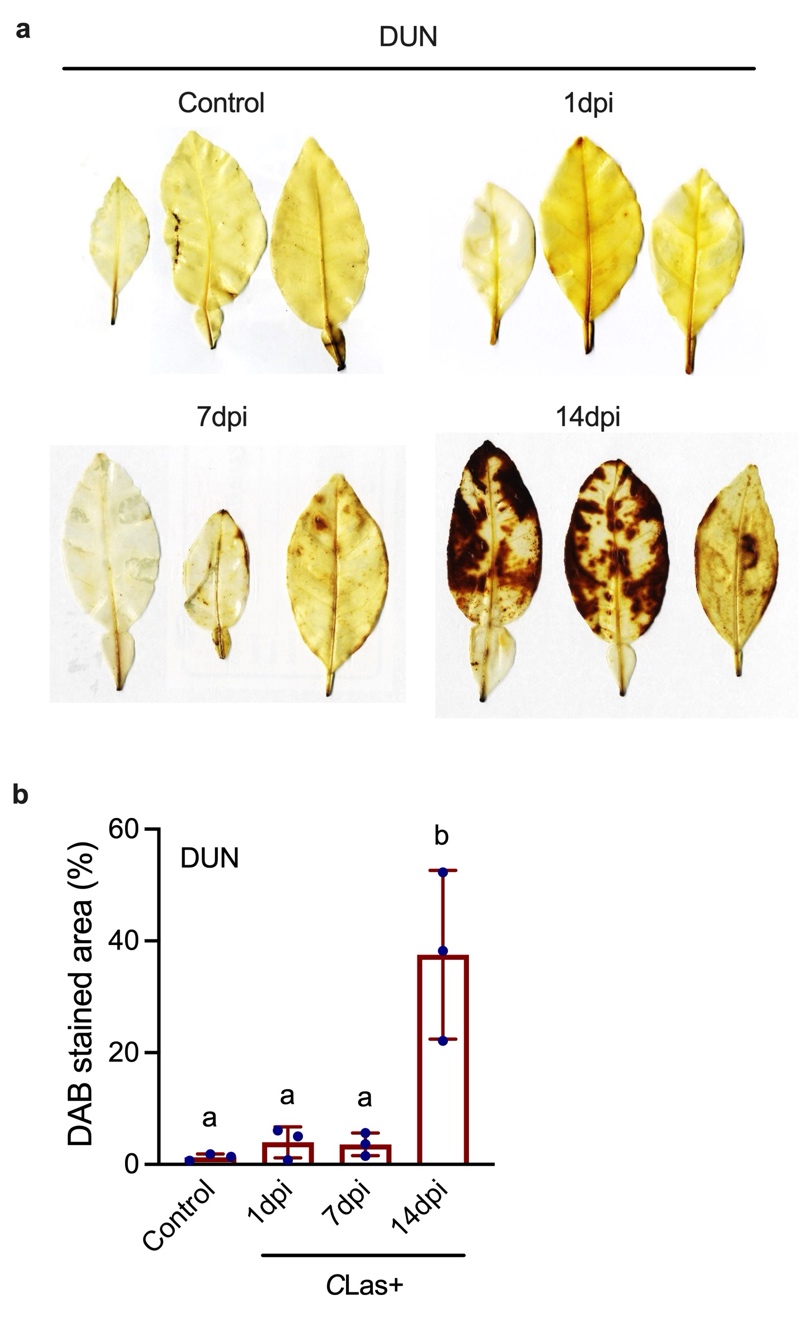

**Supplementary Figure 1** Time-course analysis of *C*Las-induced ROS accumulation in citrus leaves

**a**, ROS accumulation (brown precipitates) revealed by DAB staining in ‘Duncan’ grapefruit leaves inoculated with *C*Las-infected psyllids at 1 day post-inoculation (dpi), 7 dpi, and 14 dpi. Control: uninoculated healthy ‘Duncan’ leaves.

**b**, Percentages of leaf areas stained with DAB in the control and the ‘Duncan’ leaves inoculated with *C*Las-infected psyllids at 1, 7, and 14 dpi. Bars represent means ± SD (n = 3). Different letters denote significant differences (p < 0.05; one-way ANOVA with Tukey’s test).

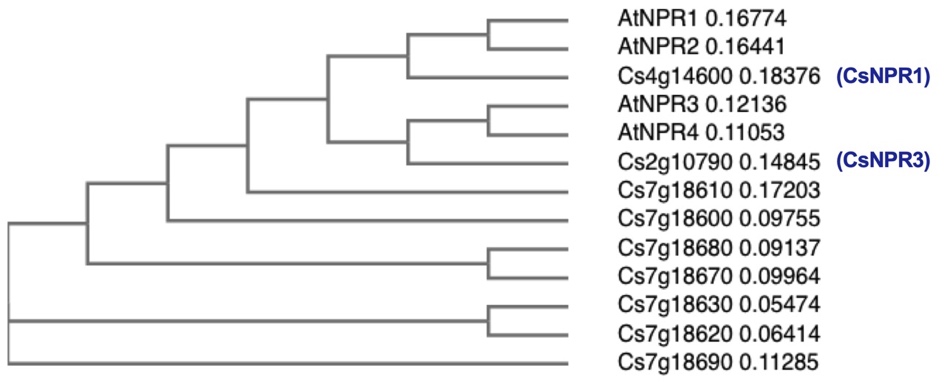

**Supplementary Figure 2** A phylogenetic tree of citrus NPR homologs, AtNPR1, AtNPR2, AtNPR3, and AtNPR4

The AtNPR1 amino acid sequence was used as the query sequence to BLAST the citrus genome database and the top nine hits plus the AtNPR proteins were used to generate the phylogenetic tree by Simple Phylogeny (https://www.ebi.ac.uk/jdispatcher/phylogeny/simple_phylogeny).

.

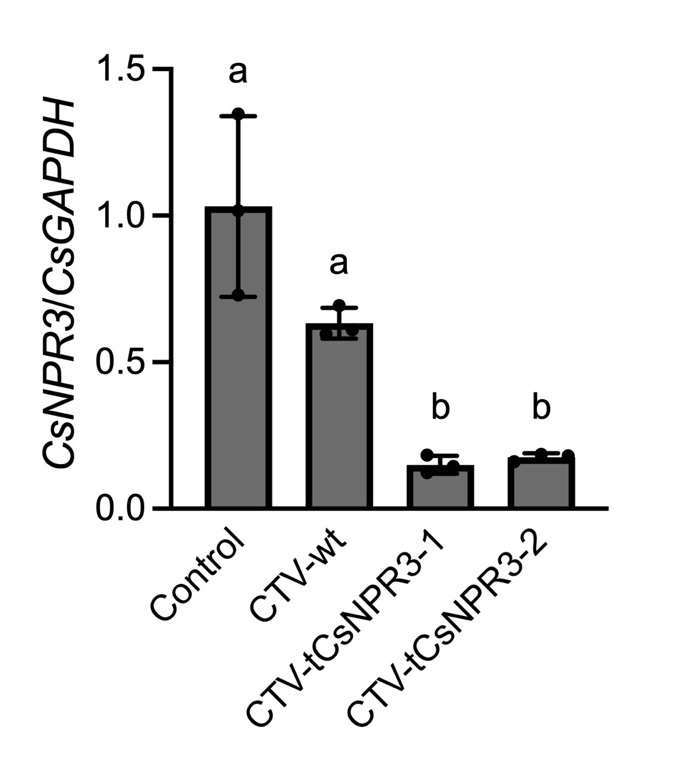

**Supplementary Figure 3** Silencing of *CsNPR3* by CTV-RNAi

Expression of *CsNPR3* in healthy (Control), CTV-wt, and CTV-tCsNPR3 *C. macrophylla* plants. Two CTV-tCsNPR3 lines were tested. Bars represent means ± SD (n = 3). Different letters denote significant differences (p < 0.05; one-way ANOVA with Tukey’s test).

**Supplementary Table 1** Primers used in this study

| Primer name | Target | Sequence (5’ to 3’) |
| --- | --- | --- |
| CQULA04F | *C*Las- 16SrDNA | TGGAGGTGTAAAAGTTGCCAAA |
| CQULA04R |  | CCAACGAAAAGATCAGATATTCCTCTA |
| *C*Las probe |  | 6FAM-ATCGTCTCGTCAAGATTGCTATCCGTGATACTAG |
| M-1636 | *CsNPR3* | CGTATGGCAAGGTTGGATATGA |
| M-1637 |  | GTTGACACCTCCATCGGAAA |
| CsCalS7-F | *CsCalS7* | GACGCCTAACCGAGTACCTGC |
| CsCalS7-R |  | GTGCAGCTGGTGATCCATCA |
| CsCalS3-F | *CsCalS3* | GGCCTCCGTTCTTACTTGCT |
| CsCalS3-R |  | ACACTCCTTGACAGCACAGG |
| CsRBOHD-F | *CsRBOHD* | CCCTCGGCTTATAAATGCAA |
| CsRBOHD-R |  | CAAAAGGCATTGAACCCAGT |
| CsGAPDH-F | *CsGAPDH* | GGAAGGTCAAGATCGGAATCAA |
| CsGAPDH-R |  | CGTCCCTCTGCAAGATGACTCT |
| StuI-tCsNPR3F | *CsNPR3* fragment | TCTAGGCCTACGTCAGCATCTGTAGAAGATTGACAA |
| PacI-tCsNPR3R |  | ACCTTAATTAACAGGTGTCTCATTTAAGTCAACCTCCCTTAA |
